# Organogenesis and Distribution of the Ocular Lymphatic Vessels in the Anterior Eye: Implication to Glaucoma Surgery Site Selection

**DOI:** 10.1101/847970

**Authors:** Yifan Wu, Young Jin Seong, Kin Li, Dongwon Choi, Eunkyung Park, George H. Daghlian, Eunson Jung, Khoa Bui, Luping Zhao, Shrimika Madhavan, Saren Daghlian, Patill Daghlian, Desmond Chin, Il-Taeg Cho, Alex K. Wong, J. Martin Heur, Sandy Zhang-Nunes, James C. Tan, Masatsugu Ema, Alex S. Huang, Young-Kwon Hong

## Abstract

Glaucoma surgeries, such as trabeculectomy, are performed to lower the intraocular pressure to reduce the risk of vision loss. The surgeries create a new passage in the eye that reroutes the aqueous humor outflow to the subconjunctival space, where the fluid is presumably absorbed by the conjunctival lymphatics. However, the current knowledge of these ocular surface lymphatics remains limited. Here, we characterized the biology and function of the ocular lymphatics using transgenic lymphatic reporter mice and rats. We found that the limbal and conjunctival lymphatic networks are progressively formed by a primary lymphatic vessel that grows out from the nasal-side medial canthus region at the time of birth. This primary lymphatic vessel immediately branches out and invades the limbus and conjunctiva, and then simultaneously encircles the cornea in a bidirectional manner. As a result, the distribution of the ocular lymphatic is significantly polarized toward the nasal side, and the limbal lymphatics are directly connected to the conjunctival lymphatics. New lymphatic spouts are mainly produced from the nasal-side limbal lymphatics, posing the nasal side of the eye more responsive to fluid drainage and inflammatory stimuli. Consistently, when a fluorescent tracer was injected, fluid clearance was much more efficient in the nasal side than the temporal side of the eyes. In comparison, blood vessels are evenly distributed on the front surface of the eyes. We found that these distinct vascular distribution patterns were also conserved in human eyes. Together, our study demonstrated that the ocular surface lymphatics are more densely present in the nasal side and uncovered the potential clinical benefits in selecting the nasal side as a surgical site for glaucoma surgeries to improve the fluid drainage.

## INTRODUCTION

The lymphatic system controls interstitial fluid homeostasis, serves as conduits for immune cell trafficking, and absorbs dietary fat and large molecules in the digestive system (1, 2). In tissue space, extravagated fluid, cells, proteins, lipids, and large molecules, collectively called lymph fluid, are reabsorbed by lymphatic capillaries, and transported back to the circulation through the pre-collectors, collecting vessels, and the thoracic duct (1, 2). Dysfunctional, damaged, and/or malformed lymphatic vessels cause abnormal interstitial fluid accumulation, which leads to interstitial pressure buildup and results in tissue swelling. Many pathological conditions, such as brain injuries, glaucoma, and lymphedema, are known to be associated with increased interstitial pressure.

Proper fluid homeostasis in the eye is critical for healthy vision and ocular health. While the lymphatics play the primary drainage function in most other tissues and organs, the fluid homeostasis in the eyes is controlled by two fluid-draining systems. In addition to the lymphatics, a specialized vascular system, called Schlemm’s canal (3–6), is also present. Schlemm’s canal is the primary route that drains aqueous humor from the anterior chamber of the eye. This specialized vascular structure is lined by a single layer of endothelial cells that displays features of both blood vascular endothelial cells (BECs) and lymphatic endothelial cells (LECs). The inner wall of Schlemm’s canal is attached to the trabecular meshwork (TM), which provides contractility for fluid drainages. Like BECs and LECs, Schlemm’s canal endothelial cells express several endothelial cell markers, such as CD31, VE-Cadherin, VEGFR-2, and vWF. We and others reported that Schlemm’s canal endothelial cells express the master regulator of lymphatics Prox1 and that Prox1 plays a crucial role in Schlemm’s canal development (3–6).

The ocular lymphatics and Schlemm’s canal share significant similarities in their organogenesis. For lymphatic vessels, a subset of venous endothelial cells upregulates Prox1 and undergoes lymphatic differentiation by upregulating LEC-signature genes and down-regulating BEC-specific genes (7, 8). These Prox1-positive lymphatic precursor cells migrate out and form the initial lymphatic vessels. In Schlemm’s canal, a subset of limbal BECs begins to express Prox1 and clusters together to form the primitive tube structure, a process termed canalogenesis (3, 4). Mature Schlemm’s canal remains connected to the blood vessels via collecting channels, which drain the aqueous humor to the systemic circulation. Functionally, both Schlemm’s canal and the ocular lymphatics play vital roles in controlling the ocular fluid homeostasis by constituting two routes of aqueous humor outflow facilities, termed conventional and non-conventional pathways respectively (9, 10). Recently, several studies have revealed previously unrecognized roles of the ocular lymphatics in eye health and disease (10–15)

Glaucoma is one of the leading causes of blindness that is characterized by optic nerve damages due to high intraocular pressures (IOP) (16, 17). Typically, the IOPs are maintained within a narrow range through a dynamic process of modulating outflow resistance through Schlemm’s canal. However, significantly increased resistance may develop primary open-angle glaucoma with ocular hypertension, which often damages the optic nerve (17). Previous reports have shown that aqueous humor flows from the ciliary processes into the anterior chamber. The aqueous humor then flows into the trabecular meshwork and the inner wall of the Schlemm’s canal, before it flows into the systemic circulation via collecting channels (18, 19). These regions are thought to contribute to aqueous humor outflow resistance that elevates the IOP (18, 19). Currently, there are numerous pharmacological and laser procedures to control high IOP in glaucoma patients. However, when these treatments fail, trabeculectomy is one of the most common surgical procedures performed to relieve ocular hypertension (17, 20). In this procedure, surgeons excise a conjunctival and scleral flap to expose Schlemm’s canal and then create a new passage between the scleral layer and the subconjunctival space of the eye. This allows aqueous humor to bypass the point of resistance at the trabecular meshwork and Schlemm’s canal (17, 20, 21). Although conjunctival lymphatics presumably absorb the drained aqueous humor, the lymphatic involvement in this process has not been undoubtedly confirmed. Therefore, it is crucial to have a better understanding of the development and function of the conjunctival lymphatic vessels and their role in maintaining ocular fluid homeostasis and normal IOP.

Previous studies have characterized limbal/corneal lymphatics in mice and humans (12, 22–33). Ecoiffier et al. have demonstrated that the limbal lymphatic vessels are unevenly distributed around the limbal circle (24). This finding provided a strong clinical significance, as the limbal lymphatics play critical roles in corneal inflammation and transplant rejection. This study has raised additional questions, especially as to when and how the ocular lymphangiogenesis occurs to establish the limbal and conjunctival lymphatic networks, as well as the functional significance of the unique ocular lymphatic distribution pattern. In this report, we performed detailed morphological studies on the organogenesis of the ocular lymphatics and Schlemm’s canal using novel lymphatic reporter transgenic animals (mouse and rat). We found that the ocular lymphatics enter the corneal limbus from the nasal side, bifurcate, and encircle the cornea, a sequential process that is very different from the development of Schlemm’s canal, which is formed by simultaneous clustering of limbal blood vessel-derived cells at the edge of the cornea (3, 4). Together, our data presented here help to advance the current understanding of the characteristic organogenesis of the ocular lymphatics and Schlemm’s canal and their roles in ocular fluid homeostasis.

## METHODS

### Animal-Related Studies

Experiments complied with the Association for Research in Vision and Ophthalmology (ARVO) Statement for the Use of Animals in Ophthalmic and Vision Research. Approval was obtained from the Institutional Animal Care and Use Committees (IACUC) at the University of Southern California (PI: YK Hong). Mice and rats were housed and raised in air-filtered clear cages in a 12-hour light/dark cycle environment and fed ad libitum. Flt1-tdsRed BAC transgenic mouse (34), Prox1-EGFP BAC transgenic mouse (35), Prox1-tdTomato BAC transgenic mouse (36), and Prox1-EGFP BAC transgenic rat (37) were previously reported. All mice were maintained in mixed backgrounds. Prox1-EGFP rats were maintained in the Sprague Dawley outbred background. For the tracer injection study, 1 µl of Bovine Serum Albumin (BSA), Alexa Fluor™ 594 conjugate (ThermoFisher Scientific) was injected into the conjunctival space of the temporal or nasal side of the left eye of anesthetized Prox1-EGFP mouse (n=6/group). After 10 minutes, mice were euthanized, and the eyes were harvested to determine the remaining amount of the tracer.

### Corneal Flat Mount Preparation

Corneal flat mounts were prepared as previously described (38). Briefly, mice and rats were euthanized by carbon dioxide asphyxiation followed by cervical dislocation or thoracic puncture, respectively. A marking hole was made on the nasal side of the eye using a syringe needle as a future directional reference. The eyes were then enucleated and fixed in 4% paraformaldehyde (PFA) at 4 °C for 2-4 hours. Fixed eyes were cut in half, and the anterior half, including the cornea, limbus, and conjunctiva, was then flattened on slides as previously described (38). All images were captured using a Leica Stereomicroscope (Leica M165 FC) or a Zeiss ApoTome Microscope (Zeiss AxioVision). Some fluorescent images were gray-scaled and then inverted to black-and-white images to increase their clarity.

### Human Tissue Studies, Histology, and Morphometric Analyses

De-identified human cornea rims were obtained after the excision of the cornea for transplantation under the approval of the Institutional Review Board (IRB) of the University of Southern California (PI: YK Hong). Tissues were fixed either in 4% PFA overnight to make frozen sections or in formalin for 48 hours to prepare paraffin sections. Immunofluorescence or immunohistochemistry assays were performed using anti-LYVE1 (R&D Systems, Minneapolis, MN) and/or anti-CD31 (Dako, Santa Clara, CA) antibodies. Vascular area and numbers were measured from > 5 representative images per group using the NIH ImageJ software. A total of three deidentified human cornea rims were collected and analyzed.

### Statistical Analyses

The two-tailed, t-test (either unpaired or paired) was used using Microsoft Excel and GraphPad PRISM6 (GraphPad Software, Inc.) to determine the statistical differences between the experimental and control groups. P-values less than 0.05 were considered statistically significant. Box-and-whisker plots were drawn using GraphPad PRISM6.

## RESULTS

### Visualization of the Ocular Surface Lymphatic Vessels and Schlemm’s Canal using transgenic reporter animals

We first characterized the distribution and molecular expression of the lymphatics present in the anterior segment of the postnatal eyes. To visualize the intact lymphatic networks, we utilized two independent bacterial artificial chromosome (BAC)-based lymphatic reporter mice that we have previously reported, namely Prox1-EGFP and Prox1-tdTomato transgenic lines (35, 36). As we and others have previously documented (3–6, 37), both the ocular lymphatics and Schlemm’s canal were readily detectable in the Prox1 promoter-driven reporter mice by their EGFP expression (Fig.1A-C). The ocular lymphatics and Schlemm’s canal demonstrated the clear morphological difference: While the limbal and conjunctival lymphatics were relatively thin and branched, Schlemm’s canal appeared to be thicker and unbranched. Also, the lymphatics are equipped with frequent luminal valves that express a high level of Prox1 (39), while Schlemm’s canal lacks equivalent valves (Fig.1B, C). Consistent with previous studies (3–6, 22), the ocular lymphatics and Schlemm’s canal commonly express Cd31 and Vegfr-3, whereas only the ocular lymphatics express Lyve1 (Fig.1D-O). The limbal lymphatics are positioned at the outer surface of the limbus, while Schlemm’s canal is located at the inner side of the limbus, near the base of the iris (Fig.1P). Moreover, Prox1 was strongly expressed in both the inner and outer walls of Schlemm’s canal.

**Figure 1.**
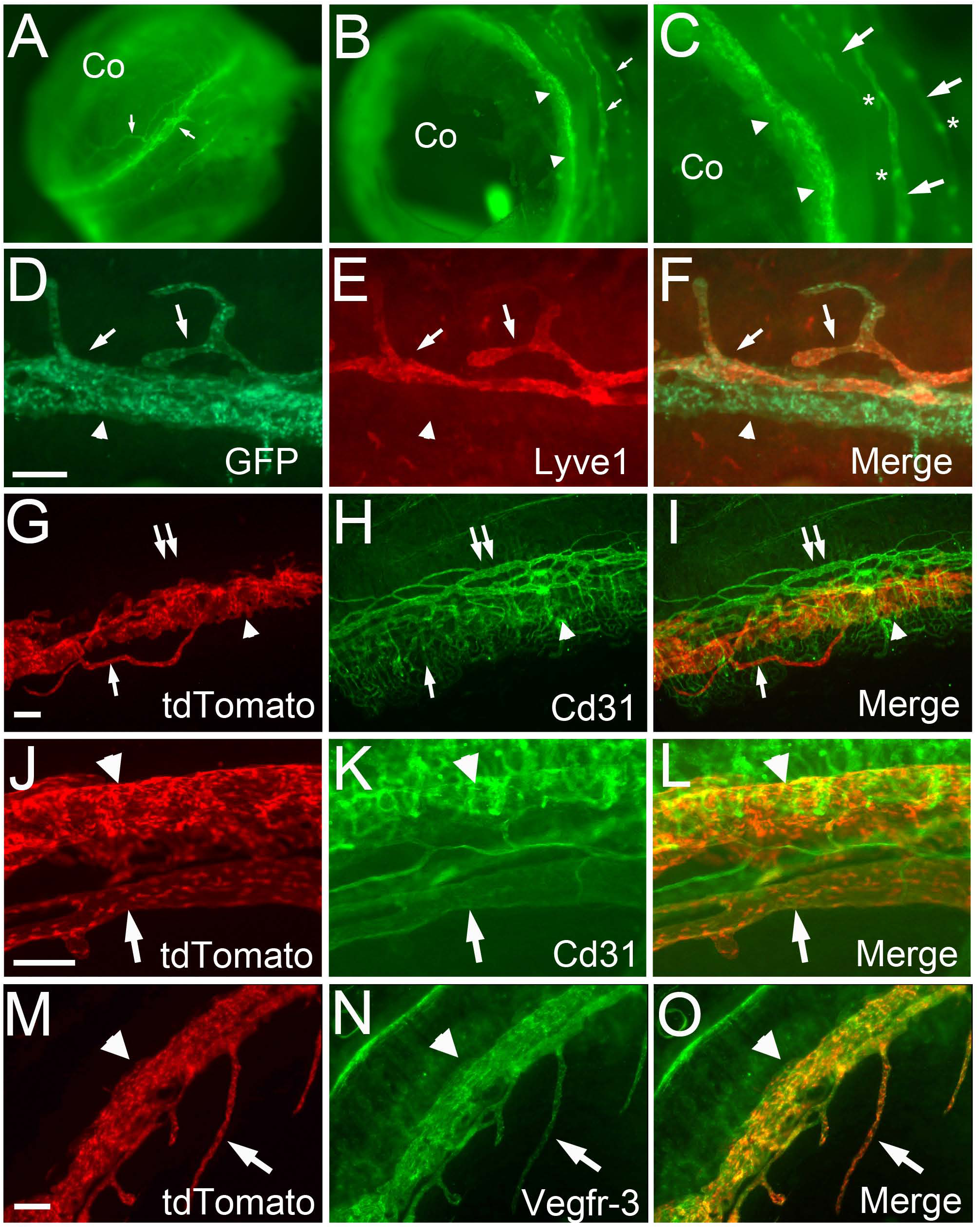
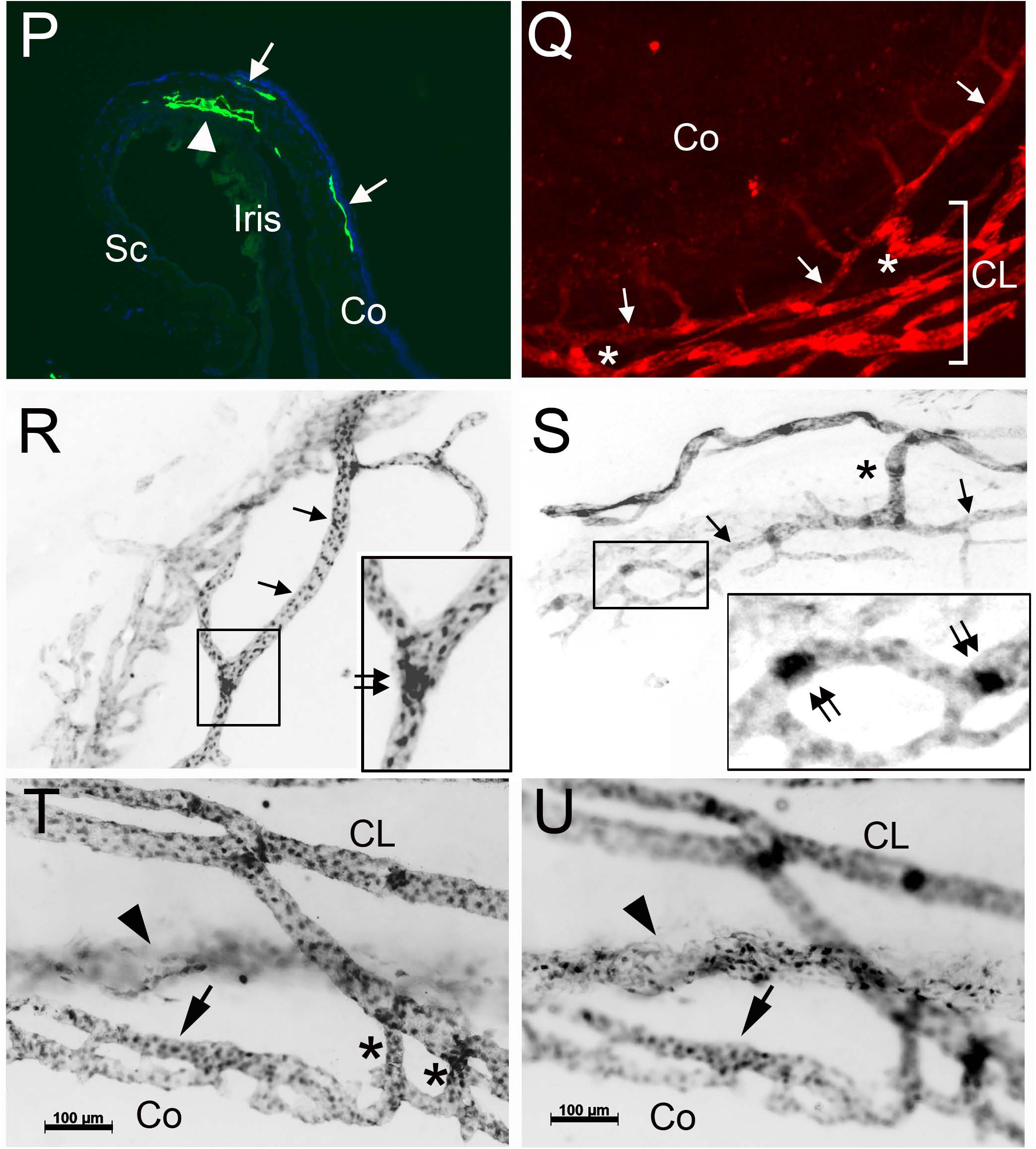
Visualization of the ocular lymphatic vessels and Schlemm’s canal. (**A-C**) Lateral views of the ocular lymphatics (arrows) and Schlemm’s canal (arrowheads) in the anterior segment of the eye of Prox1-EGFP transgenic mouse. The limbus area in panel B is enlarged in panel C, where lymphatic valves (asterisks) are marked with stronger EGFP (Prox1) expression than luminal lymphatic endothelial cells. (**D-F**) Lyve1 whole-mount staining (red) showing the limbal lymphatics and Schlemm’s canal in the eye of Prox1-EGFP mouse. While EGFP (Prox1) is expressed in both Schlemm’s canal (arrowhead) and the limbal lymphatics (arrows), Lyve1 is detected only in the limbal lymphatics, not in Schlemm’s canal. (**G-L**) Anti-Cd31 staining (green) of the limbal blood and lymphatic vessels and Schlemm’s canal in the eyes of Prox1-tdTomato reporter mouse. Schlemm’s canal (arrowhead) and the limbal lymphatics (arrow), but not limbal blood vessels (double arrows), are selectively shown by Prox1-tdTomato reporter (red). (**M-O**) Anti-Vegfr-3 staining (green) in the eyes of Prox1-tdTomato reporter mouse. Vegfr-3 is expressed in both the limbal lymphatics (arrow) and Schlemm’s canal (arrowhead). (**P**) Expression of the reporter (EGFP) in the inner and outer walls of Schlemm’s canal (arrowhead) as well as the limbal lymphatics (arrow) in a frozen cross-section of the eye of Prox1-EGFP mouse. (**Q-S**) The limbal lymphatics (arrows) are directly connected to the conjunctival lymphatics (CL) and equipped with lymphatic valves. The connections are marked with asterisk (*) in panels, Q and S, and the lymphatic valves (double arrows) are shown in panels, R and S, where the boxed areas are enlarged in the insets. (**T,U**) High magnification images, focusing on either the lymphatics (T) or Schlemm’s canal (U), in the same location of the eyes of Prox1-EGFP rat (postnatal day 5). The limbal lymphatics (arrow), Schlemm’s canal (arrowhead), and conjunctival lymphatics (CL) are visualized, and the limbal-to-conjunctival lymphatic connection is marked with asterisks. Co, Cornea; Sc, Sclera. Scale bars: 100 µm. More than three eyes from the reporter animals (both genders, 6-10 weeks old) were used for each experiment and one representative image is shown for each panel.

It has not been reported whether the limbal lymphatics harbor the luminal valves, a hallmark of collecting lymphatics, and how they extend beyond the limbus area. Toward these questions, we next performed further studies on the limbal lymphatics using two different rodent species, Prox1-tdTomato mice (36) and Prox1-EGFP transgenic rat (37). Indeed, we found that the limbal lymphatics are directly connected to the conjunctival lymphatics through multiple short lymphatics present along the corneal edge (Fig.1Q-U). We also detected the Prox1^−high^ luminal valves in mature limbal lymphatics in adults (Fig.1R,S), suggesting non-capillary characters of limbal lymphatics (2, 40). Together, the ocular lymphatics and Schlemm’s canal display distinct anatomic and molecular characteristics.

### *In Vivo* Detection of the Ocular Lymphatic Vessel Connection to the Extraocular Tissues

We next set out to perform in-depth morphological analyses on the ocular lymphatics using Prox1-tdTomato mice (36) and Prox1-EGFP transgenic rat (37). The entire heads of the reporter animals were fixed with paraformaldehyde to preserve the structures of the ocular and surrounding tissues. We found that the limbus and conjunctival tissues, but not the cornea, were richly endowed with the lymphatic networks (**Fig.2**). Importantly, the lymphatics were more densely distributed in the nasal side than the temporal side, and these ocular lymphatic networks appeared to exit toward the medial canthus of the eye in both species. Enlarged images revealed that a strand of the large-caliber lymphatic vessel comes into the nasal-side conjunctiva from the medial canthus, and branches out to form the conjunctival lymphatic networks. Together, the lymphatic reporter mice and rats commonly revealed the abundant lymphatic networks in the conjunctiva, which appeared to originate from the medial canthus area in the nasal side.

**Figure 2.**
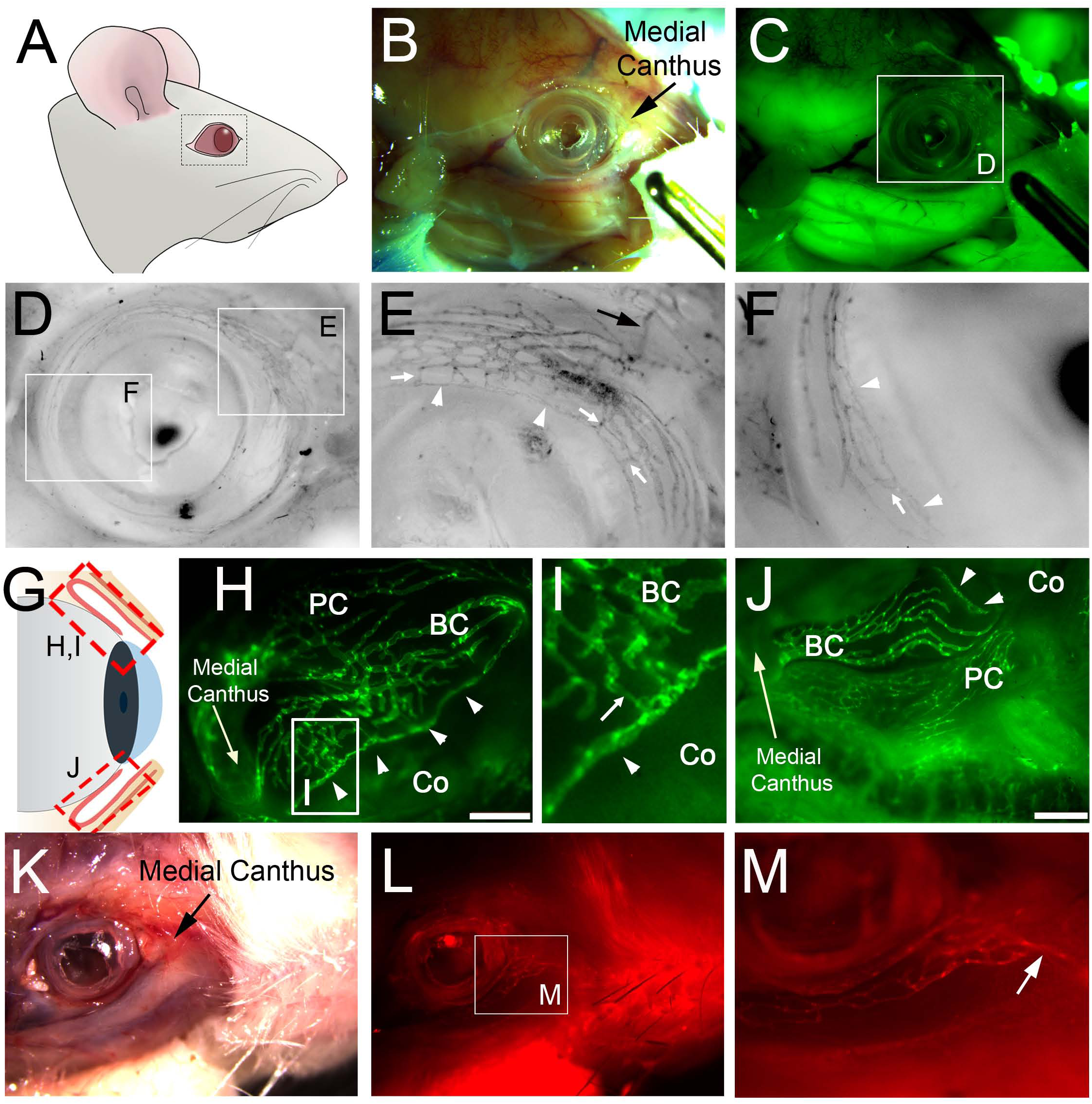
*In situ* visualization of the lymphatic vessels in the ocular and surrounding tissues. **(A-F)** The entire head of Prox1-EGFP rat was fixed and partially dissected to reveal the ocular lymphatics surrounding the ocular tissue. (**A**) Diagram of the rat head showing the region of interest. (**B, C**) Bright-field and fluorescence images of the partially skinned head of Prox1-EGFP rat. (**D**) Inverted and gray-scaled image of the ocular lymphatics with two boxed areas that are enlarged in panels (**E**) and (**F**). The limbal lymphatics (white arrowheads) are directly attached to the conjunctival lymphatics through multiple short branches (white arrows). (**G-J**) Dense lymphatic networks in the conjunctiva. (**G**) Simplified diagram showing the superior (**H, I**) and inferior (**J**) conjunctiva in the nasal side. (**I**) The boxed area in panel (**H**) was enlarged. Arrow points the connection between the conjunctival lymphatics and the limbal lymphatics (marked with arrowheads). PC, palpebral conjunctiva; BC, bulbar conjunctiva; Co, cornea. Scale bars, 1 mm. (**K-M**) The head of Prox1-tdTomato mouse was fixed and partially skinned to reveal the lymphatics surrounding the ocular tissue. Arrow in panels (**E**) and (**M**) points to a main lymphatic vessel that exits the ocular tissue and extends toward the medial canthus. More than three rats and four mice (both genders) were used for each experiment with comparable results.

### Stepwise Growth of the Ocular Lymphatics from the Nasal Side in Mouse and Rat

The lymphatic distribution shown above prompted us to investigate further how the ocular lymphatics postnatally develop and form the dense networks in the conjunctiva and limbus. To address this question, we enucleated the developing eyes of the lymphatic reporter newborn pups and adults, and prepared corneal flat mounts to visualize the postnatal ocular lymphangiogenesis. We found that a nascent lymphatic vessel, originating from the nasal side of the eye, grows into the conjunctiva area, then bifurcates and sprouts both clockwise and counter-clockwise, and eventually covers the limbal and conjunctiva area during the first two weeks after birth (Fig.3A-F). The limbal and conjunctival lymphatics concurrently encircled the cornea in parallel and made frequent connections among them through short perpendicular branches as shown above (Fig.1.Q-U). In comparison, Schlemm’s canal independently developed without any connections to the ocular lymphatics, and became visible from postnatal day 6-8 (arrows, Fig.3 G,H). Quantitative analyses revealed that about 3-4 times more ocular (limbal and conjunctival) lymphatic vessels were formed in the nasal side, compared to the temporal side, and this polarized lymphatic distribution is maintained in adults (Fig.3I,J). Intriguingly, we noted that all ocular lymphatics seem to grow out from one lymphatic sac, which may serve as the root of the ocular lymphatics (Fig.3K). It is possible that this sac potentially funnels the lymph fluid taken by all ocular lymphatics to the regional lymph nodes and eventually to the circulation. Thus, the identity, character, and function of this enigmatic lymphatic sac need further investigation. We also confirmed this pattern of nasal-side predilection of the ocular lymphatic development using a different species, Prox1-EGFP lymphatic reporter rats (37). Indeed, we observed a comparable pattern of ocular lymphangiogenesis in rats: the limbal and conjunctival lymphatics in newborn rat pups (P2) migrated from the nasal side and bifurcated to encircle the cornea (Fig.3L). Notably, Schlemm’s canal did not yet formed at this point. At P9, while the limbal and conjunctival lymphatics are in the process of encircling the cornea, Schlemm’s canal developed evenly throughout the corneal edge at this time point (Fig.3M.N). It is interesting to find that the ocular lymphatics originated from one side (nasal) of the eye and grew toward the other side (temporal), encircling the cornea over two weeks, whereas Schlemm’s canal uniformly established over a few days only around P5-P8 (3–6). Together, this study provides detailed information about the developmental timing and morphogenesis of the ocular lymphatic networks.

**Figure 3.**
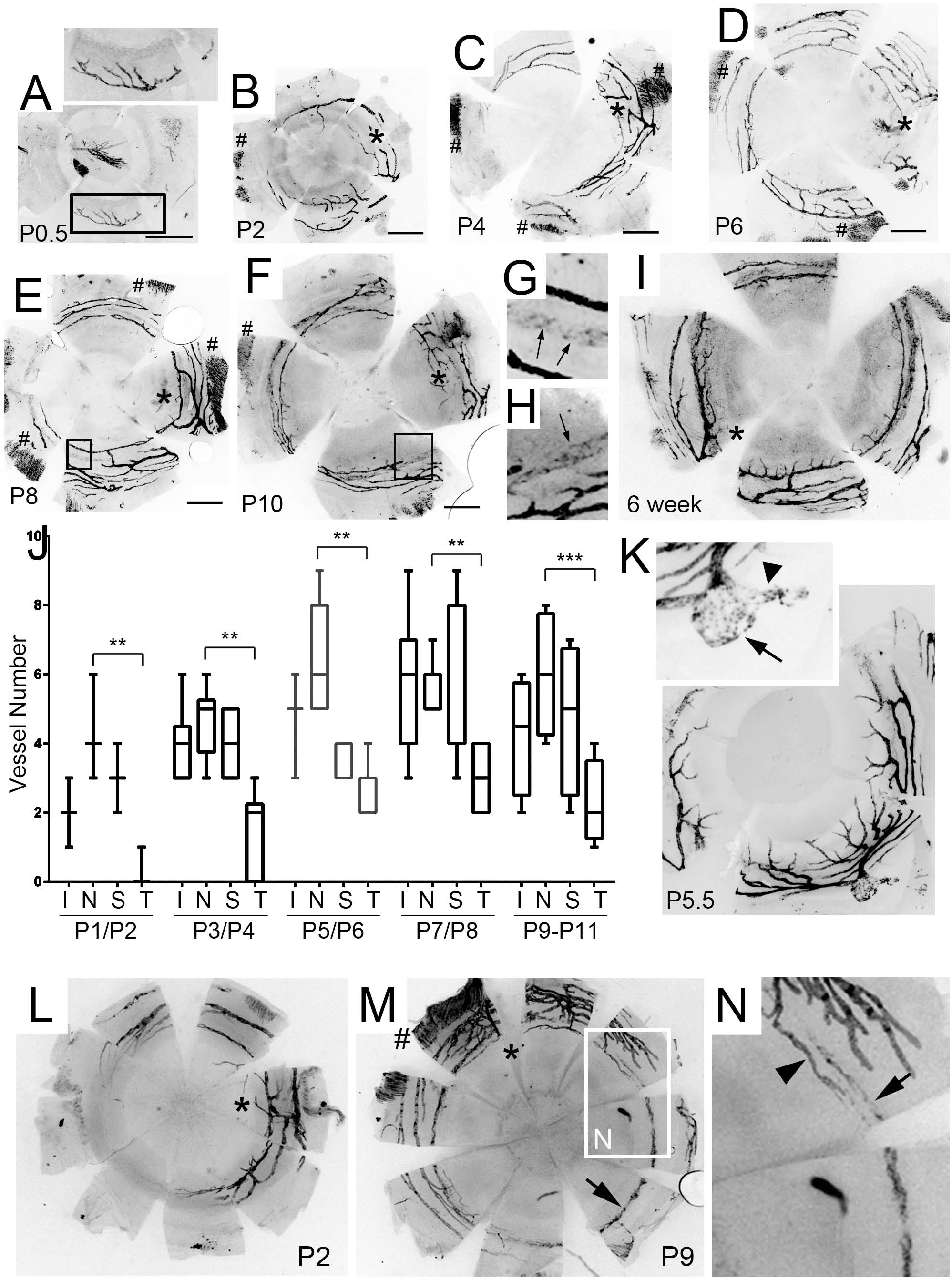
Postnatal Development of the Ocular Lymphatics Originating from the Nasal Side in Mouse. (**A-I**) Images showing the sequential lymphatic growth in the conjunctiva and limbus. The eyes from Prox1 reporter pups at postnatal (P) days 0.5, 2, 4, 6, 8, and 10, and 6-weeks old adult were enucleated, and cornea flat mounts were prepared. Asterisks (*) point the nasal side of the eye and pound (#) signs label the ocular muscles for the expression of Prox1 in muscles (35, 37). Note that the ocular lymphatics originate from the nasal side, and grow into the limbus (the inner circle) and conjunctiva (outer circles). Formation of Schlemm’s canal (arrows) is evident after P6-8 and the boxed area in panels D and E are enlarged in panels G and H, respectively. All scale bars, 0.5 mm. 3-7 pups (both genders) per time point were used for each experiment and one representative image is shown. (**J**) A box plot showing the quantitative analyses of lymphatic vessels encircling the cornea in the Inferior (I), Nasal (N), Superior (S), and Temporal (T) quadrants of the eyes at the indicated postnatal days P1/P2 (n=3), P3/P4 (n=6), P5/P6 (n=7), P7/P8 (n=7), and P9-P11 (n=4). (**K**) Discovery of a presumptive initial lymph sac in the eye of the Prox1 reporter pup (P5.5). In the enlarged inset, an arrow points the presumptive lymph sac, which may serve as a root for all the ocular lymphatics and funnel the lymph fluid to the regional lymph nodes. Arrowhead marks a connection of this sac to the downstream collecting lymphatics. (**L-N**) Cornea flat mounts prepared from Prox1-EGFP reporter rats at P2 (**L**) and P9 (**M**) showing that the ocular lymphatics in rats also originate from the nasal side (asterisks). The boxed area in panel (**M**) is enlarged in panel (**N**) to show the limbal lymphatic vessels (arrowhead) growing around the cornea edge, and Schlemm’s canal (arrow) forming nearby. #, the ocular muscles. Statistical significance (p-value) was calculated by unpaired (for panel J) or paired (for panel N) *t*-test (two-tailed), **, *p* < 0.01; ***, *p* < 0.001.

### More Efficient Conjunctival Drainage in the Nasal Side than the Temporal Side

We next asked whether fluid drainage is more efficient in the nasal side than the temporal side due to higher lymphatic density in the nasal side conjunctiva. To compare the drainage of the two sides, we injected a fluorescent tracer (Albumin-Red, MW= 66kDa) into the nasal or temporal side of the eyes of Prox1-EGFP mice. Ten minutes post-injection, the mice were euthanized, and the cornea flat preps were prepared to compare the remaining amount of the tracer on each side. Indeed, the tracer injected into the nasal side of the conjunctiva was completely drained, but a significant amount of the tracer injected into the temporal side largely remained in the conjunctiva (**Fig.4**). Together, this study demonstrates differential drainage rates between the temporal and nasal sides, consistent with the polarized distribution of the conjunctiva lymphatics.

**Figure 4.**
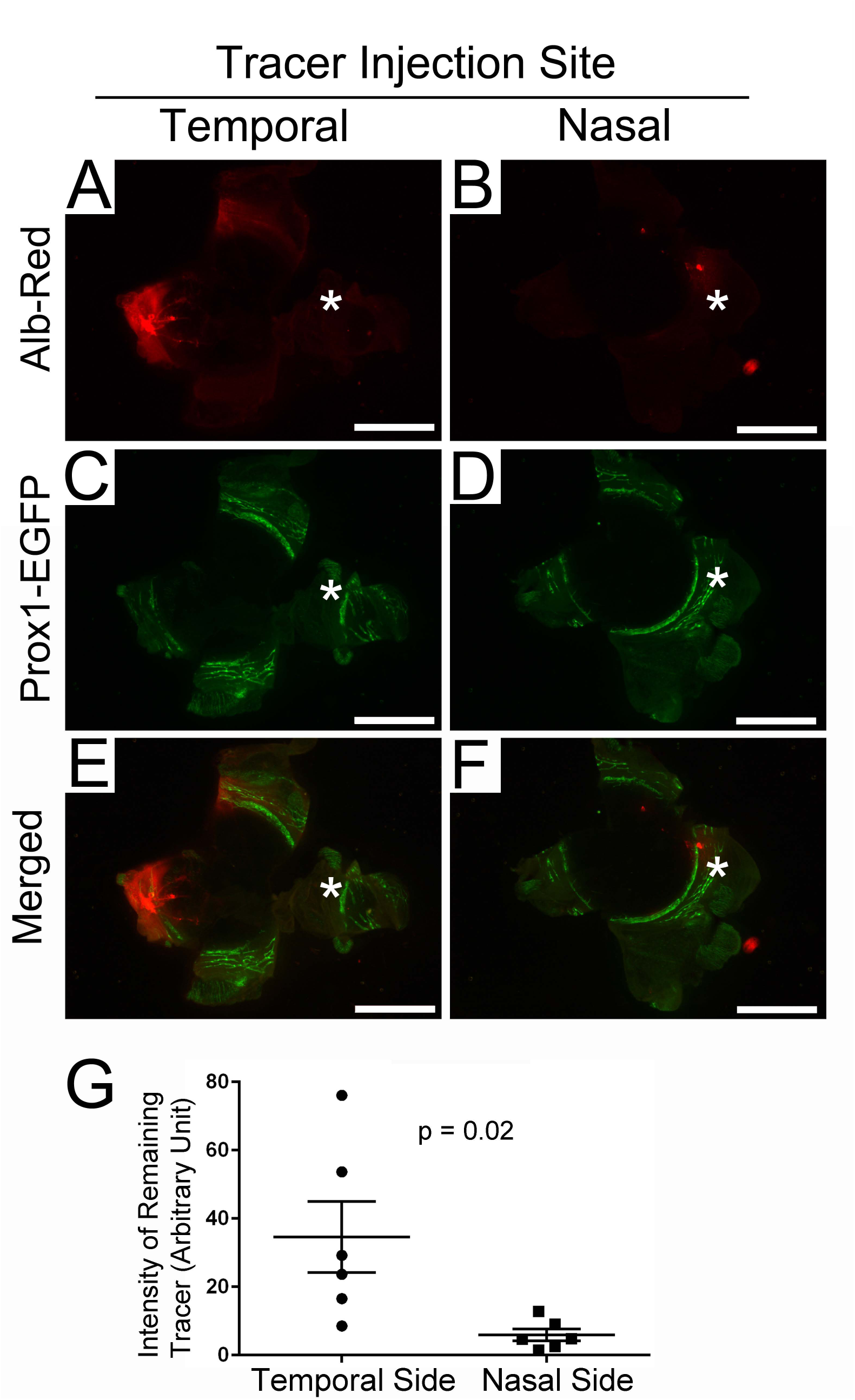
Faster fluid drainage in the nasal side compared to the temporal side. A red fluorescence tracer (Albumin-Red, 1 µl) was injected into the conjunctiva of the temporal side (**A, C, E**), or nasal side (**B, D, F**) of the left eye of individual Prox1-EGFP mouse. The same injection pressure was applied for all injections by a microinjector. After 10 minutes, the animals were euthanized, and their eyes were harvested. The remaining tracer in the cornea prep was quantified by measuring the intensity using NIH ImageJ software (**G**). Asterisks mark the nasal side. Alb-Red, albumin-Red; scale bars, 2 mm. Statistical significance (p-value) was calculated by unpaired *t*-test (two-tailed) on mice (both genders). Six eyes were used for each group, and 12 mice were used for the experiment.

### Spontaneous Pathological Ocular Lymphangiogenesis in Neonatal Pups

We noted that the limbal lymphatics give out multiple fine sprouts toward the cornea and that these thin lymphatic sprouts form mainly from the nasal-side limbal lymphatics, but rarely from the temporal-side limbal lymphatics (Fig.5A-C). These short blunt-ended lymphatic sprouts from the nasal-side limbal lymphatics seemed to be poised to respond to potential infection, playing a role in the ocular immune response. Consistent with these patterns, we detected a case of spontaneous pathological corneal lymphangiogenesis at postnatal P7, where an array of the limbal lymphatic sprouts was growing toward the center of the avascular cornea (Fig.5D). Remarkably, this extensive pathological lymphangiogenesis mainly occurred from the nasal side (Fig.5E), while Schlemm’s canal remained unresponsive and did not generate any sprouts (Fig.5F). Moreover, we also detected perinatal pathological lymphangiogenesis at an earlier time point. In an eye of a newborn mouse (P0.5), pathological lymphangiogenesis occurred before physiological lymphatic development (Fig.5G). It was interesting to note that both developmental and pathological lymphangiogenesis were concurrently occurring. Notably, their vascular growth patterns were significantly different: while developmental lymphangiogenesis generated an organized, stepwise and fractal lymphatic growth from the nasal side of the eye (Fig.5H), the pathological lymphangiogenesis initiated from a separate vascular root and hastily formed a non-fractal lymphatic network (Fig.5I).

**Figure 5.**
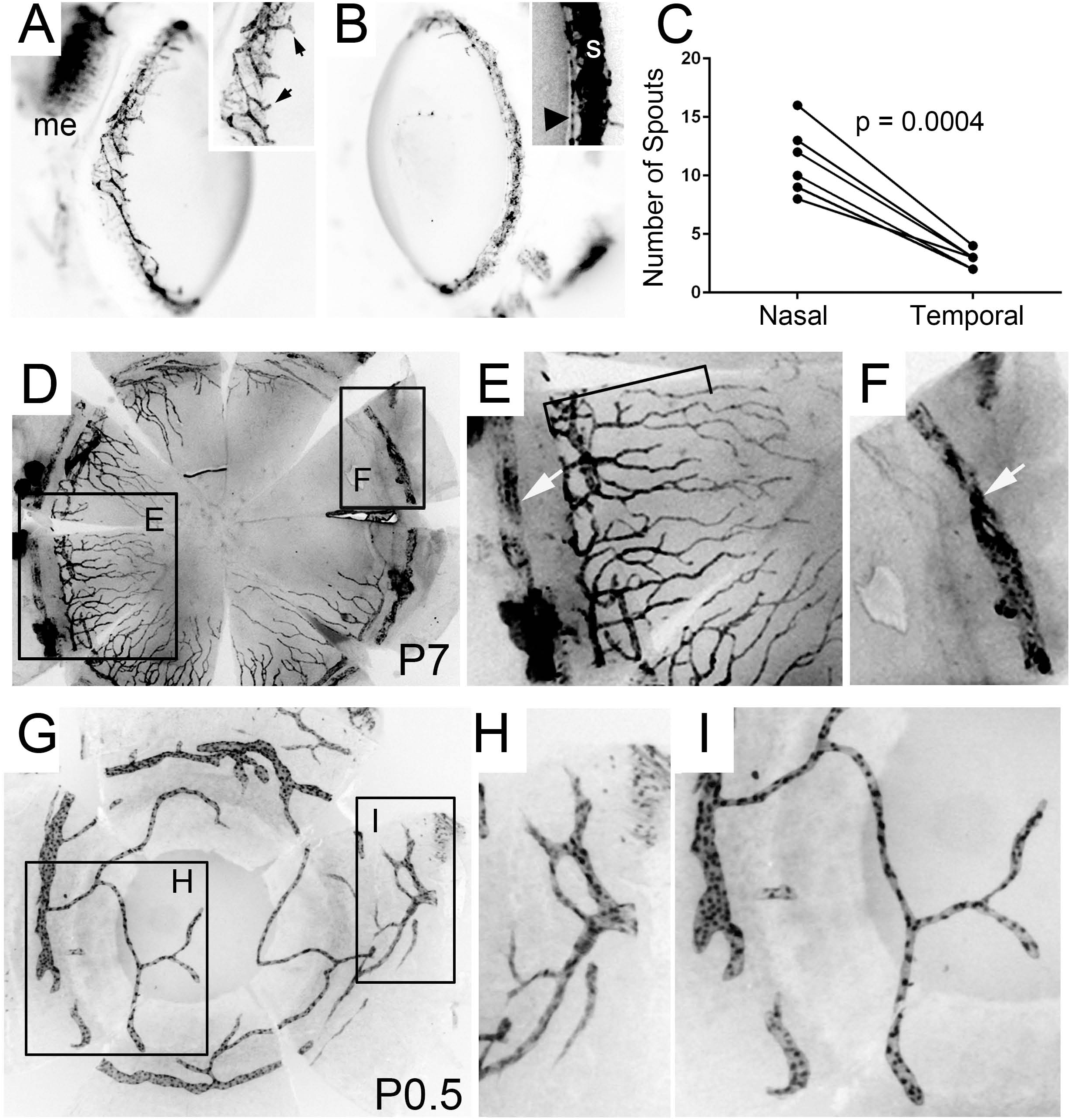
Pathological Ocular Lymphangiogenesis. (**A-C**) Short lymphatic sprouts are found in the nasal-side limbal lymphatics. Lateral views of the nasal side (**A**) and temporal side (**B**) of the eye of healthy Prox1-tdTomato mouse (8 weeks old) demonstrate the predominant presence of limbal lymphatic sprouts in the nasal side. A part of the nasal and temporal lymphatics was enlarged and shown in the insets. Black arrows, sprouts from the limbal lymphatic vessel; Black arrowhead, the limbal lymphatic vessel; S, Schlemm’s canal; me, medial rectus muscle. (**C**) The number of limbal lymphatic sprouts in the nasal *vs*. temporal quadrant was quantified (n=6, both genders). (**D-F**) Extensive corneal lymphangiogenesis in the postnatal eye of Prox1-EGFP rat pup (P7) due to unknown causes. White arrows point Schlemm’s canal, and a bracket marks active corneal lymphangiogenesis. Boxed areas in panel (**D**) are enlarged in panels (**E**) and (**F**), respectively. (**G-I**) Concurrent occurrence of developmental and pathological lymphangiogenesis detected in the eye of Prox1-EGFP neonate mouse (P0.5). Boxed areas in panel (**G**) are enlarged in panels (**H**) and (**I**), respectively.

### Comparative Analyses of the Postnatal Ocular Angiogenesis and Lymphangiogenesis

As the limbal and conjunctival area is also heavily endowed with blood vessels (42, 43), we next performed comparative analyses of the angiogenesis and lymphangiogenesis in the anterior segment of the eyes. We first stained the anterior region of the eyes from Prox1-tdTomato pups (P4) against Cd31, a pan-endothelial cell marker that is highly expressed in blood vessels but only weakly in lymphatic vessels (44). While lymphatic vessels were still growing and encircling the cornea, blood vessels had already covered the sclera, limbus, and iris regions at this developmental stage (Fig.6A-C). Moreover, whereas a main lymphatic trunk originated from the nasal side, multiple blood vessels, including long and anterior ciliary arteries, had already entered the limbus area. This is consistent with the previous report that the limbus vasculature primarily derives from the anterior ciliary arteries and their branches form the vascular arcades to perfuse the limbal conjunctiva and peripheral cornea (45). We next employed the Flt1-tdsRed reporter mouse line (34), which marks blood vascular endothelial cells with a red color-emitting tdsRed protein. Flt1-tdsRed reporter was mated with the Prox1-EGFP line to produce double transgenic mice (Flt1-tdsRed/Prox1-EGFP) to visualize both blood and lymphatic vessels simultaneously. At P1, the limbus and conjunctiva were fully covered with blood vessels, as previously shown (46), while the lymphatics began to invade the limbal area (Fig.6D-G). At P3 and P5, while blood vessels were evenly distributed, the lymphatic vessels were still in the progress of encircling the cornea toward the temporal side (Fig.6H-O). Together, our study demonstrates that blood vessel development precedes lymphatic vessel development in the anterior part of the eyes and that both vasculatures independently develop without any significant spatiotemporal overlaps.

**Figure 6.**
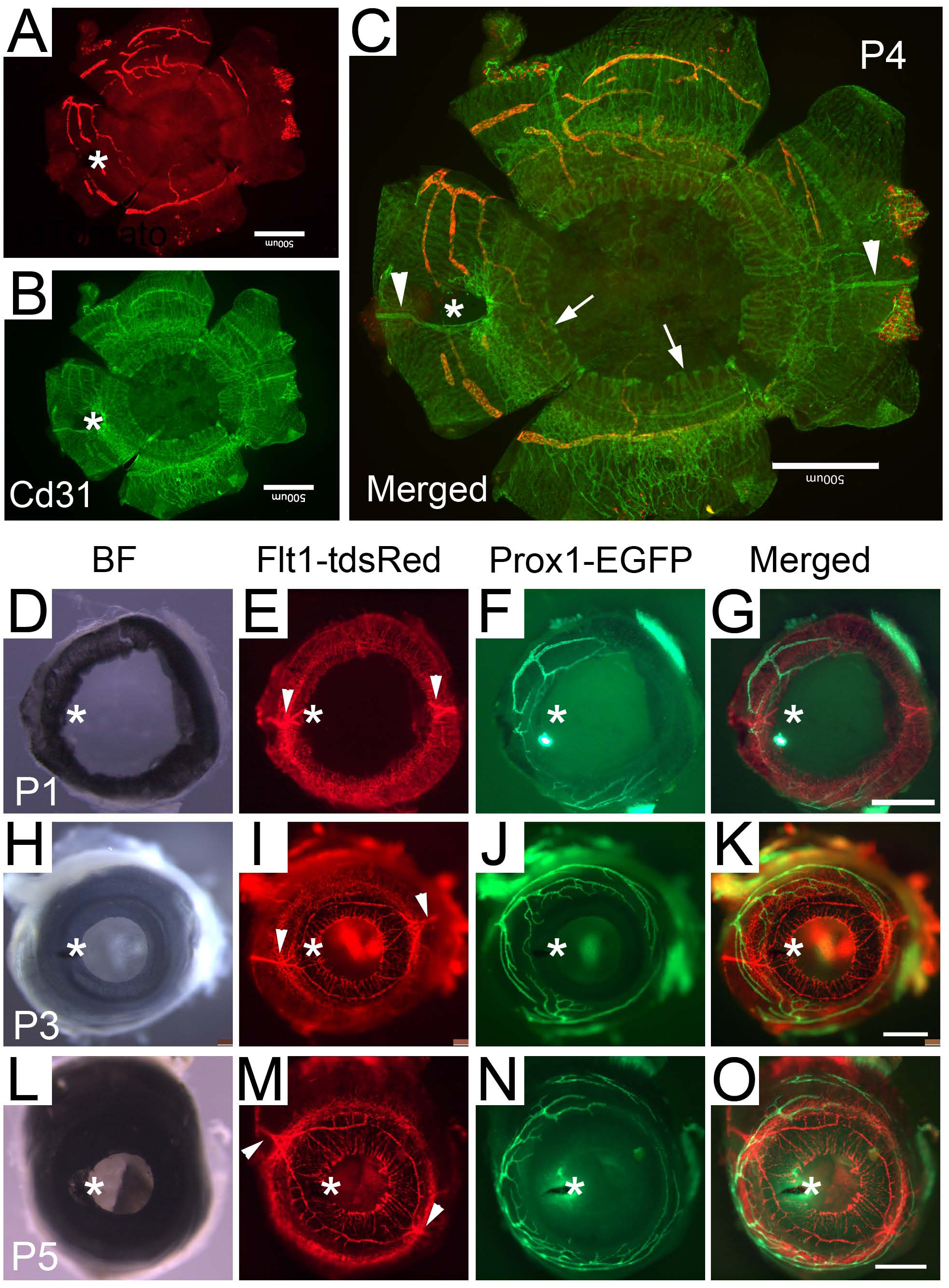
Postnatal Development of Blood and Lymphatic Vessels in the Anterior Eye. (**A-C**) A flat corneal mount of the eye of Prox1-tdTomato reporter pup (P4) was stained for Cd31. Lymphatic vessels (tdTomato) (**A**) and Cd31-stained vessels (green) (**B**) revealed distinct and independent development of the two vasculatures in the anterior segment of the eye. (**C**) Merged image. (**D-O**) Front views of the anterior eyes of Flt1-tdsRed/Prox1-EGFP double transgenic pups revealed characteristic development of the ocular blood and lymphatic vessels at P1 (**D-G**), P3 (**H-K**), and P5 (**L-O**). Asterisk marks the nasal side. Arrowheads indicate long ciliary arteries, and arrows point to the iris. Scale bars, 500 µm.

### Polarized Distribution of Lymphatic Vessels in Human Conjunctiva

We next asked whether the polarized distribution of the conjunctiva lymphatics could also be seen in humans. To address this question, we obtained human cornea rims after corneal transplantation and performed histological analyses focused on the peri-limbal conjunctiva. All examined cornea rims retained the intact corneal-scleral junction and a skirt of the bulbar conjunctiva. To assess the vascular distribution, we equally segmented the cornea rims into six fragments, prepared their cross-sections, and measured their vascular density. Indeed, anti-LYVE1 staining revealed a polarized distribution of lymphatic vessels (*i.e.*, vessel number & density) in the conjunctival area, whereas anti-CD31 staining revealed largely even vascular distribution throughout all sections (Fig.7). This finding in human eyes was consistent with the observations that we made in rodent eyes. Nonetheless, we were unable to precisely and confidently identify the orientation (nasal vs. temporal) of the corneal rims due to the limited anatomic information available for the tissues. In conclusion, human eyes also display a polarized lymphatic distribution, like those of rodents.

**Fig.7.**
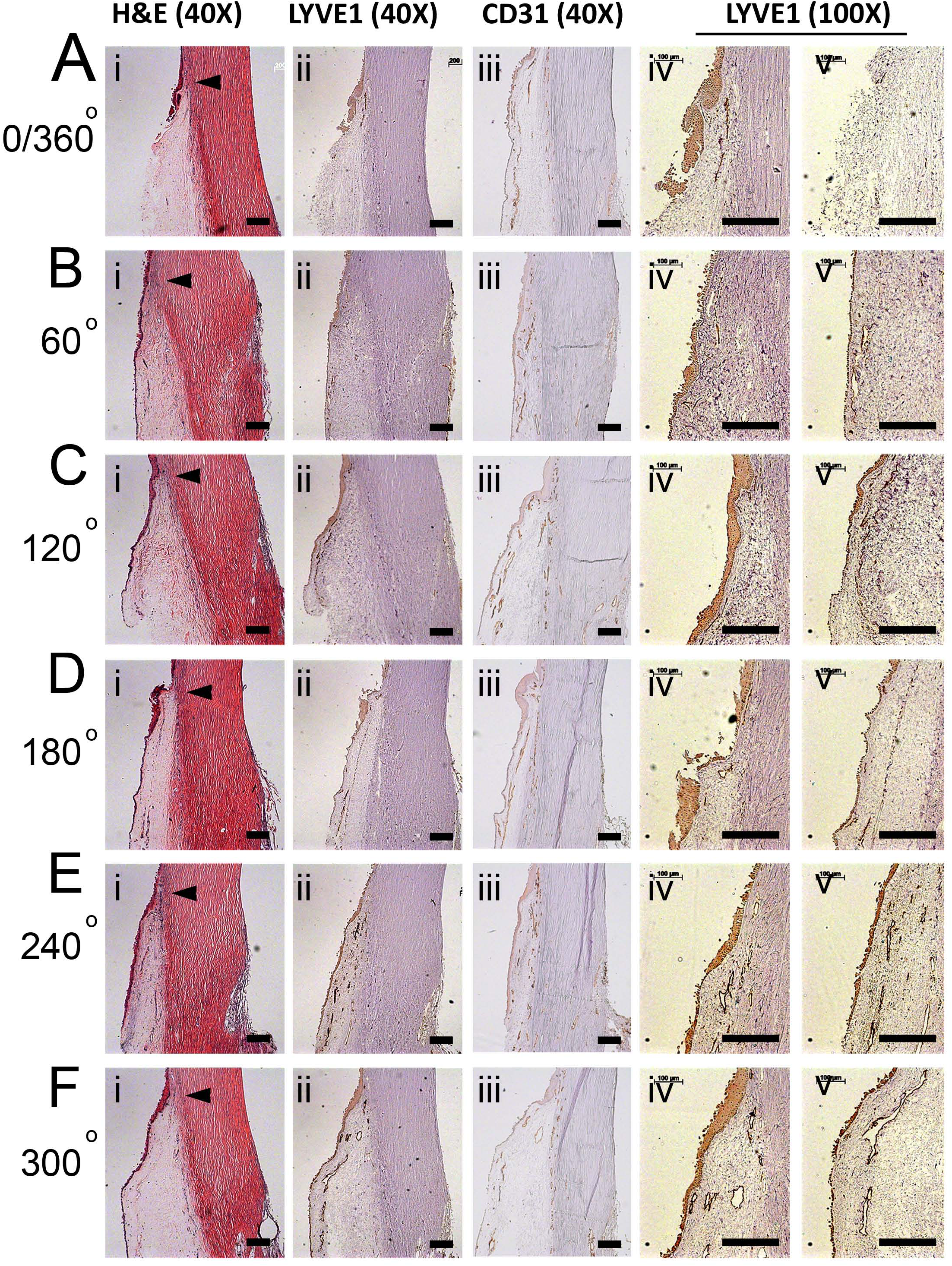
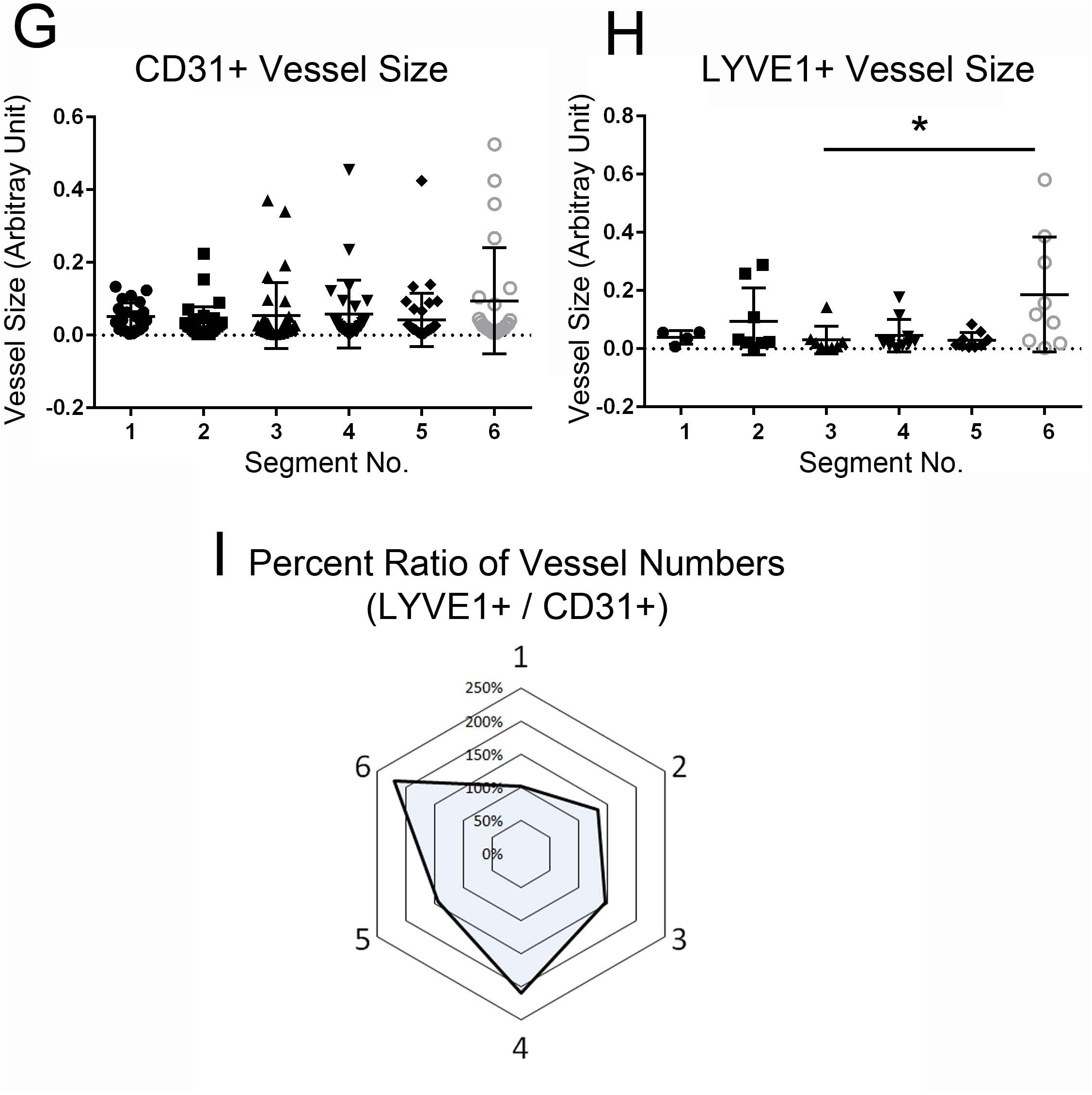
Distribution of Conjunctival Blood and Lymphatic Vessels in the Anterior Part of the Human Eyes. Cornea rim tissues, remaining after excision of the cornea for transplantation, were equally divided into 6 segments with incisions 60-degrees apart from each other (**A-F**), paraffin-embedded, and cross-sectioned for immunohistochemistry for H&E (**i**), LYVE1 (**ii**), and CD31 (**iii**) images taken at 40X magnification, and for LYVE1 images (**iv**, **v**) taken at 100X magnification. Arrowheads point to the cornea-conjunctiva junctions. Scale bars, 200 µm. (**G-I**) Morphometric analyses were performed to quantify the vessel size (area) for CD31-positive vessels (**G**) and LYVE1-positive vessels (**H**) in each segment. Segment No.1 is the 0/360° segment in panel (**A**). (**I**) A percent ratio of lymphatic vs. blood vessel number was calculated as follows. Segment numbers are shown at the corners of the hexagon. In each segment, the total LYVE1 vessel number was divided by the total CD31 vessel number, and then the resulting number was converted to a percentage by setting the number of Segment No.1 to 100%. A total of three human eye rims were analyzed with consistent results.

## DISCUSSION

In this study, we investigated the development and function of the ocular lymphatics and Schlemm’s canal. Our results highlighted significant differences in morphological structures, development timeline, and vessel distribution patterns between the ocular lymphangiogenesis and Schlemm’s canal formation using three independent lymphatic reporter animals. Especially, we illustrated the stepwise process of the ocular lymphatic network formation (Fig. 8). The ocular lymphangiogenesis begins with nascent lymphatic vessels emerging from the nasal side of the developing eye. These rapidly growing lymphatics sprout and bifurcate, and soon encircle the cornea in both clockwise and counter-clockwise. During this process, the limbal and conjunctival lymphatics concurrently grow toward the temporal side of the eye, making frequent connections among them, and eventually cover the entire conjunctival area. As a result, the nasal-side is more densely enriched with the lymphatics, compared to the temporal side. In addition, more lymphatic sprouts were also produced by the limbal lymphatics in the nasal-side. The polarized ocular lymphatics raised a possibility that fluid drainage and immune surveillance may be more efficient in the nasal-side than the temporal side of the eye. Indeed, our results found that the nasal side was more efficient in tracer drainages and more responsive to postnatal infection compared to the temporal side of the eye. In comparison, the blood vessels in the anterior surface of the eyes did not display this kind of polarized vascular pattern. Importantly, this nasal-side predilection of the ocular lymphatics was also conserved in human eyes.

**Fig.8.**
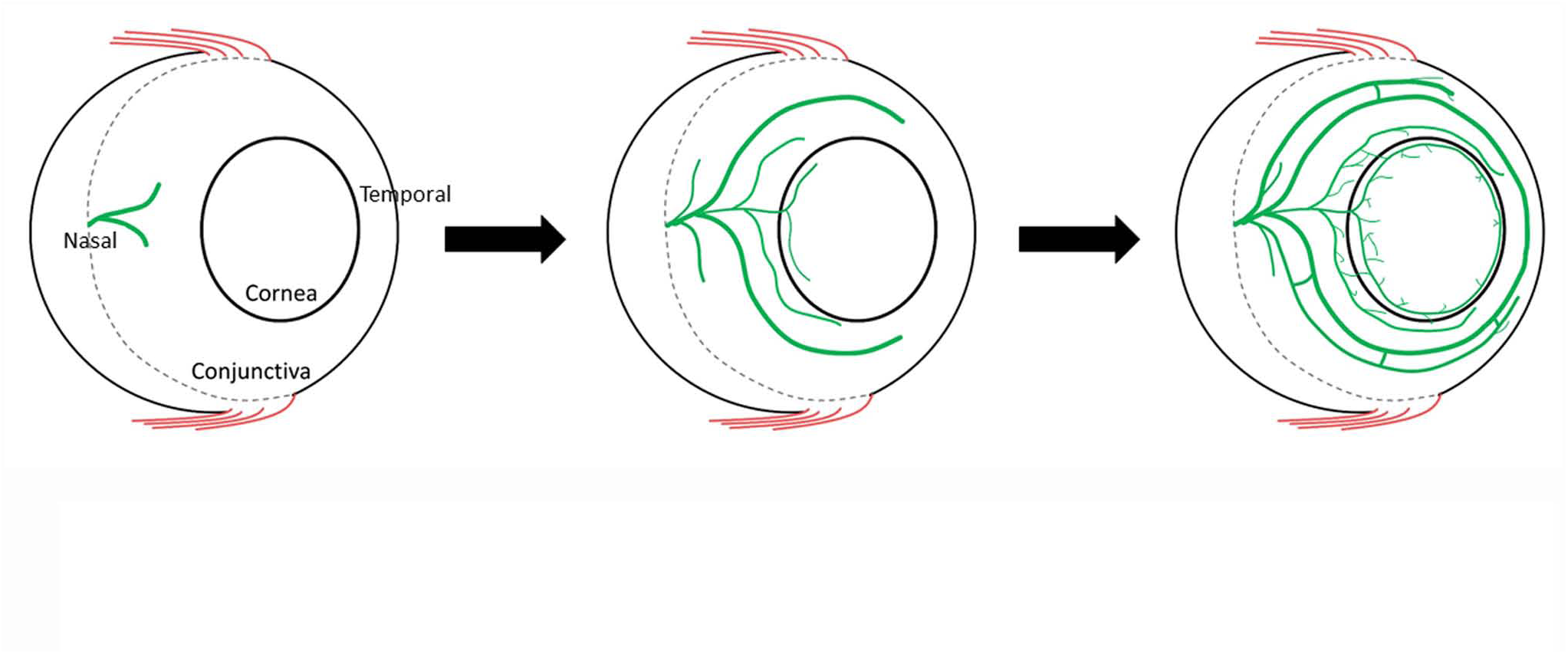
Stepwise Development of the Ocular Lymphatic Plexus. A simplified diagram illustrating the stepwise development of the postnatal limbal and conjunctival lymphatics.

The outcomes of our study provide an important implication to glaucoma surgeries, which aim to lower the intraocular pressure (IOP) and reduce the risk of glaucoma. Glaucoma surgeries generally create new passages that allow rerouting of the aqueous humor out of the anterior chamber. As a result, rerouted fluid accumulates and forms a small reservoir, also called as bleb, at the subconjunctival space until the fluid is presumably drained by the lymphatics nearby. It has been proposed that proper post-trabeculectomy fluid drainage and IOP management are heavily dependent on the function of the ocular lymphatic system and its association with the formed blebs (47, 48). Based on a tracer introduction, trabeculectomy blebs showing a clear lymphatic outflow have been correlated with better IOP lowering results (47, 48). An interesting implication raised by these studies was that Mitomycin C, routinely treated on the surgery site to limit tissue scarring that could block the newly created passages, could inhibit the growth of these draining lymphatics and potentially negatively affect the outcome of the surgery. In agreement with this notion, a previous study found that failed blebs previously treated with Mitomycin C resulted in decreased lymphatic and blood vessel density (20). Thus, healthy lymphatics may play a major role in reducing elevated IOP and fluid clearance in bleb-forming glaucoma surgeries. As such, the lymphatics are known to reside in the conjunctiva and postulated to play a key role in draining the bleb fluid (49, 50). Currently, trabeculectomy blebs are often placed in the superior or temporal quadrants to minimize potential post-surgery infections and to be covered by the eyelid. Since our study uncovered the nasal-side predilection of the conjunctival lymphatics, a nasal or superior-nasal placement of the bleb may be other options that are potentially beneficial to access these lymphatics for better outflow drainage and infection management.

On the other hand, understanding the polarized vascular pattern of the ocular lymphatics would be clinically important for drug delivery. In this sense, the outcome of our study also provides an important implication for ocular drug delivery. Subconjunctival space often acts as an important reservoir for iatrogenically injected subconjunctival medicines, such as antibiotics and steroids, for ocular disease treatment through transscleral delivery. Drugs injected into the subconjunctival space also form another type of bleb. The injected drug will be either delivered through the sclera to the targets or wastefully clearly by the conjunctival lymphatics. Thus, the presence of abundant lymphatics nearby the bleb could negatively impact the delivery of the drugs, as these drugs could be rapidly drained and cleared by the conjunctival lymphatic outflow. Together, avoiding the lymphatic-rich region of the eye by injecting drugs on the temporal side may enhance the drug’s longevity and subsequent permeability through the sclera into the eye, as opposed to leaving the eye via drainage by the lymphatics.

In conclusion, our work demonstrates the spatiotemporal development and polarized distribution of the ocular surface lymphatics to the nasal side of the eye in mice and rats. Consistent with these data, our human data supports the nasal predilection of the ocular surface lymphatics. This may have clinical relevance for glaucoma treatment and drug delivery, which can deliver a significant impact on all aspects of eye care. Future work can be guided by developmental results and directed at identifying pharmacological/molecular tools to manipulate lymphatic presence (either greater or lesser) to aid in the treatment of eye diseases.

## ACKNOWLEDGMENTS

This study was supported by the National Institutes of Health (Grant Numbers: EY026260, HL121036, HL141857, DE027891, DK114645 to YH; K08EY024674 to ASH; K08 HL132110 to AKW) and by American Cancer Center (#IRG-16-181-57 to DC).

